## Supplemental Figures for "The lncRNA Malat1 Inhibits miR-15/16 to Enhance Cytotoxic T Cell Activation and Memory Cell Formation"

### Supplemental Data & Figures

#### Figure S1. *Malat1$^{scr}$* allele does not disrupt other miRNA families

TargetScan predicted binding sites for highly expressed microRNA families in T cells that contained at least one HITS-CLIP read in both WT and Malat1$^{scr/scr}$ CD8$^{+}$ T cells were compared for depth of Ago2 HITS-CLIP reads. First, reads at the predicted seed site were normalized by total Ago2 HITS-CLIP reads in a given 3' UTR. To best visualize all sites, logit transforms of these values are plotted. Paired t-test performed to determine significance. Blue line indicates the identity line. Data for each genotype is from combined libraries of n = 2 biological replicates.

(A) Predicted sites for let-7-5p
(B) Predicted sites for miR-21-5p
(C) Predicted sites for miR-101-3p
(D) Predicted sites for miR-142-3p

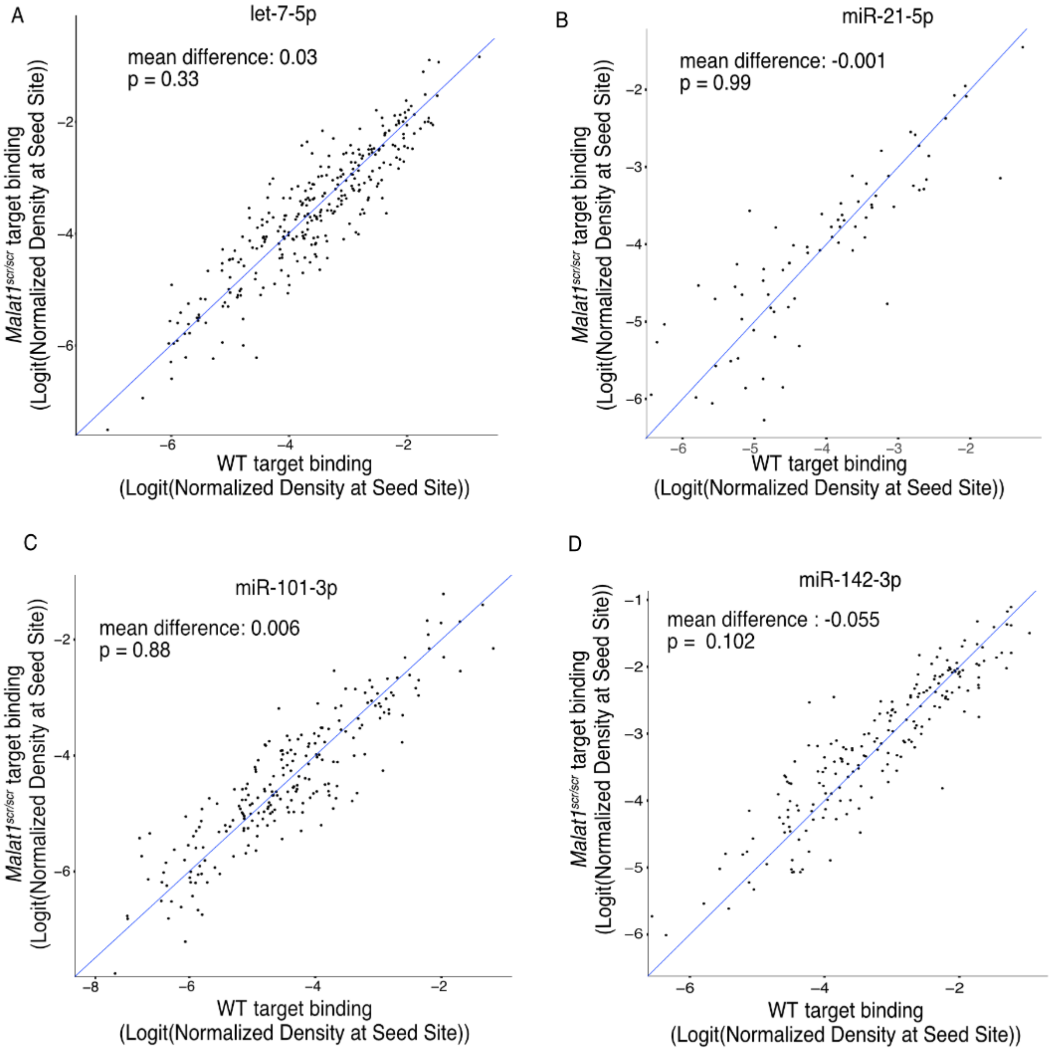
A
let-7-5p
mean difference: 0.03
p = 0.33
Malat1 scr/scr target binding
(Logit(Normalized Density at Seed Site))
WT target binding
(Logit(Normalized Density at Seed Site))
B
miR-21-5p
mean difference: -0.001
p = 0.99
Malat1 scr/scr target binding
(Logit(Normalized Density at Seed Site))
WT target binding
(Logit(Normalized Density at Seed Site))
C
miR-101-3p
mean difference: 0.006
p = 0.88
Malat1 scr/scr target binding
(Logit(Normalized Density at Seed Site))
WT target binding
(Logit(Normalized Density at Seed Site))
D
miR-142-3p
mean difference : -0.055
p = 0.102
Malat1 scr/scr target binding
(Logit(Normalized Density at Seed Site))
WT target binding
(Logit(Normalized Density at Seed Site))

### Figure S2. AHC and gene expression analyses nominate direct miR-15/16 targets involved in growth signaling pathways

(A) Heat map indicating differential gene expression between WT and *Malat1*$^{scr/scr}$ cells, and between miR-15/16$^{fl/fl}$ cells and miR-15/16$^{\Delta/\Delta}$ cells in RNA sequencing analyses.
(B-C) Histograms of AHC mapping of Ago2 binding in the 3' UTR of *Pik3r1* (B) and *Mapk8* (C) in WT (black), miR-15/16$^{\Delta/\Delta}$ (red) and *Malat1*$^{scr/scr}$ (blue) cells. As in Figure 3, AHC libraries were generated from CD8$^{+}$ T cells isolated from spleens and cultured for 5 days (combined libraries from n = 2 for each genotype). Grey bars indicate the peaks containing TargetScan predicted miR-15/16 binding sites.

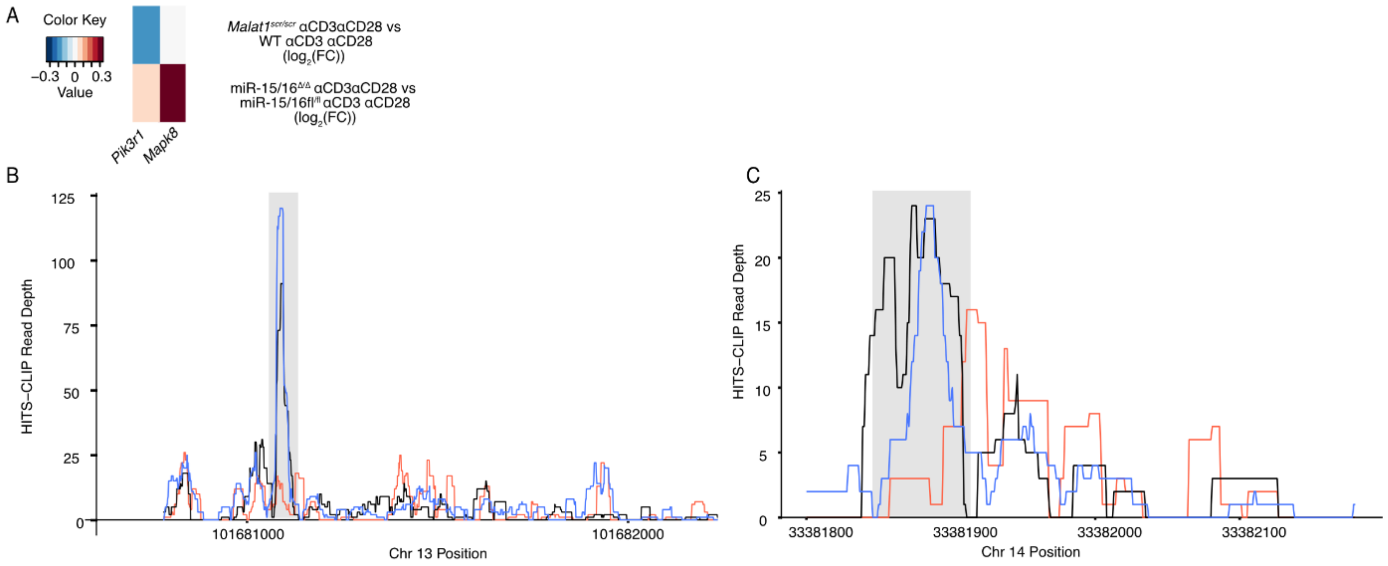

A
Color Key
−0.3 0 0.3
Value
Malat1scr/scr αCD3αCD28 vs
WT αCD3 αCD28
(log2(FC))
miR-15/16Δ/Δ αCD3αCD28 vs
miR-15/16fl/fl αCD3 αCD28
(log2(FC))
Pik3r1
Mapk8
B
HITS-CLIP Read Depth
125
100
75
50
25
0
101681000
101682000
Chr 13 Position
C
HITS-CLIP Read Depth
25
20
15
10
5
0
33381800
33381900
33382000
33382100
Chr 14 Position

### Figure S3. CD28 responsive genes are induced by αCD28 stimulation in all genotypes tested

Cumulative density plots comparing expression of CD28 responsive gene set defined as genes from (Martínez-Llordella et al., 2013) with αCD3αCD28 vs αCD3 $\log_2$(FC) > 1.5 and adjusted p value < 0.001. Kolmogorov-Smirnov test used to determine significant differences in the distributions of target and non-target genes. αCD3 used at 1 μg/mL, and α CD28 used at 1 μg/mL. Data are from a single experiment with n=6 for each genotype and stimulation condition combination.

(A) Comparison of CD28 responsive genes in WT cells
(B) Comparison of CD28 responsive genes in *Malat1*$^{scr/scr}$ cells
(C) Comparison of CD28 responsive genes in miR-15/16$^{fl/fl}$ cells
(D) Comparison of CD28 responsive genes in miR-15/16$^{\Delta/\Delta}$ cells

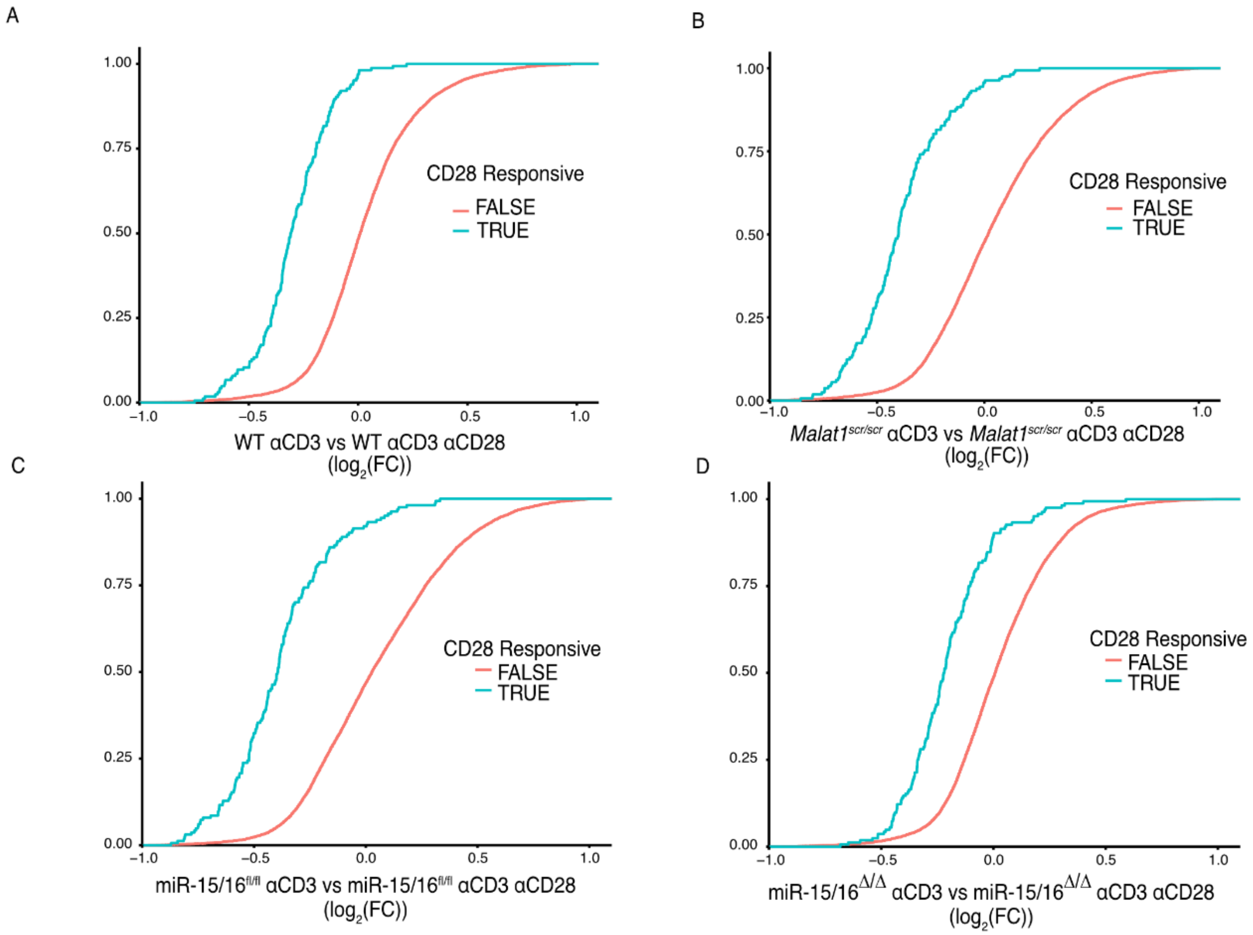
A
1.00
0.75
0.50
0.25
0.00
−1.0
−0.5
0.0
0.5
1.0
CD28 Responsive
FALSE
TRUE
WT αCD3 vs WT αCD3 αCD28
(log2(FC))
B
Malat1scr/scr αCD3 vs Malat1scr/scr αCD3 αCD28
(log2(FC))
CD28 Responsive
FALSE
TRUE
C
miR-15/16fl/fl αCD3 vs miR-15/16fl/fl αCD3 αCD28
(log2(FC))
CD28 Responsive
FALSE
TRUE
D
miR-15/16Δ/Δ αCD3 vs miR-15/16Δ/Δ αCD3 αCD28
(log2(FC))
CD28 Responsive
FALSE
TRUE

### Figure S4. Malat1 Regulates Memory Formation in Unchallenged Poly Clonal Animals

Cells were isolated from the spleens of young, age-matched, naive mice and analyzed by flow cytometry for CD44 and CD62L to delineate naive, effector memory, and central memory cells. Results shown are gated on $CD8^{+}$ $CD5^{+}$ lymphocytes. Data for *Malat1*$^{scr/scr}$ and WT cells are from 3 independent experiments. Data for miR-15/16$^{fl/fl}$ and miR-15/16$^{\Delta/\Delta}$ cells are from 2 independent experiments. Statistics displayed determined by unpaired t-test between Malat1$^{scr/scr}$ and WT cells or between miR-15/16$^{fl/fl}$ and miR-15/16$^{\Delta/\Delta}$ cells (*, $p<0.05$; **, $p<0.01$)
(A) Representative flow cytometry plots of CD44 and CD62L for each genotype assayed. Percentages shown are of CD8+ population.
(B) Quantification of naive cells ($CD62L^{+}$ $CD44^{-}$)
(C) Quantification of central memory cells ($CD62L^{+}$ $CD44^{+}$)
(D) Quantification of effector memory cells ($CD62L^{-}$ $CD44^{+}$)

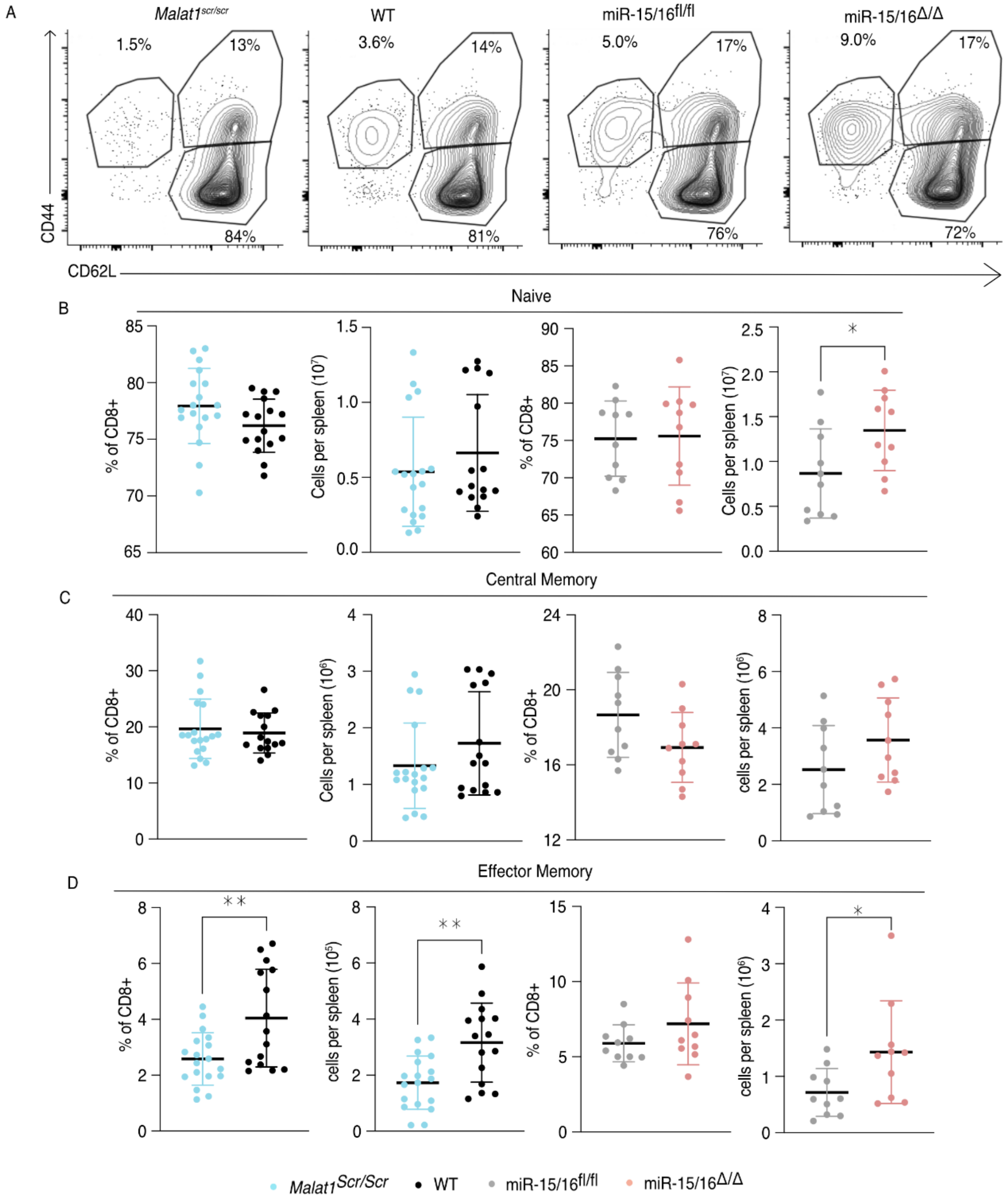
A
Malat1scr/scr
WT
miR-15/16fl/fl
miR-15/16Δ/Δ
1.5%
13%
84%
3.6%
14%
81%
5.0%
17%
76%
9.0%
17%
72%
CD44
CD62L
B
Naive
% of CD8+
Cells per spleen (10^7)
*
C
Central Memory
% of CD8+
Cells per spleen (10^6)
cells per spleen (10^6)
D
Effector Memory
% of CD8+
cells per spleen (10^5)
cells per spleen (10^6)
**
**
*
Malat1Scr/Scr
WT
miR-15/16fl/fl
miR-15/16Δ/Δ

### Figure S5. Malat1 is epistatic to miR-15/16 in the regulation memory cell expansion following LCMV infection

Polyclonal mice of the miR-15/16$^{fl/fl}$, miR-15/16$^{\Delta/\Delta}$, and *Malat1*$^{scr/scr}$;miR-15/16$^{\Delta/\Delta}$ genotypes were directly infected i.p. with 5*$10^5$ p.f.u. I.p. lcmv armstrong. Antigen specific responses were tracked in the blood and spleen using the GP33 tetramer at day 7 and day 31 post infection.

(A) Representative flow cytometry plots illustrating the gating on antigen specific cells using CD44 and GP33

(B) Quantification of GP33+ cells in the blood at day 7

(C) Quantification of GP33+ cells in the spleen at day 31 by absolute numbers and percent of the CD8+ T cell population

(D) Representative flow cytometry plots of KLRG1 and CD127 expression at day 31 in the antigen specific cell population

(E) Representative flow cytometry plots of CD27 and CD43 expression at day 31 in the antigen specific cell population

(F) Quantification of KLRG1$^{+}$ antigen specific cells in the blood at day 7 by relative percentage and absolute numbers

(G) Quantification of KLRG1+ antigen specific cells in the spleen at day 31 by relative percentage and absolute numbers

(H) Quantification of CD43+ CD27+ memory cells in the spleen at day 31 by relative percentage and absolute numbers

(I) Quantification of CD43- CD27- t-Tem cells in the spleen at day 31 by relative percentage and absolute numbers

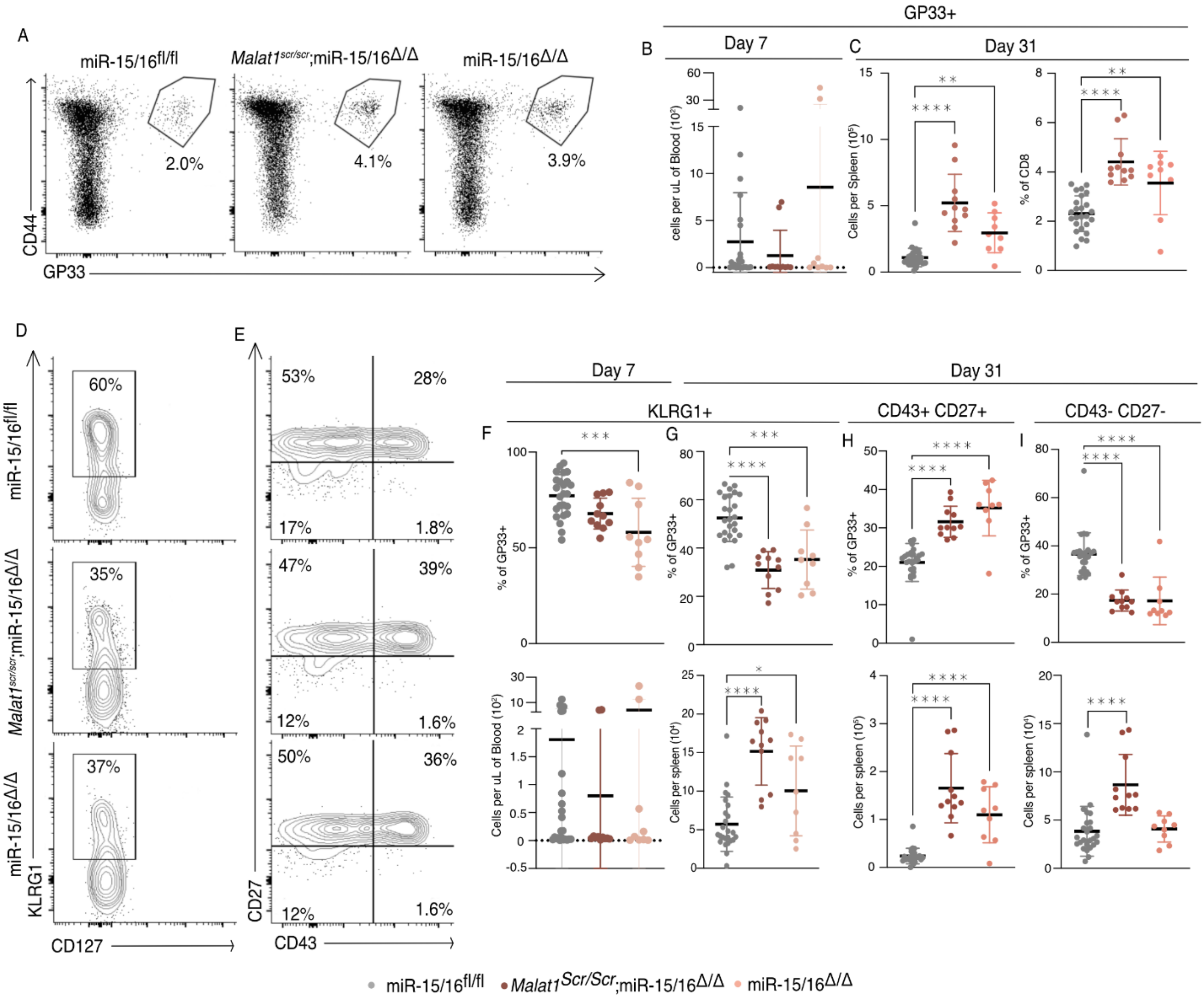
A
miR-15/16fl/fl
Malat1scr/scr;miR-15/16Δ/Δ
miR-15/16Δ/Δ
2.0%
4.1%
3.9%
CD44
GP33
GP33+
B
Day 7
C
Day 31
cells per uL of Blood (10²)
Cells per Spleen (10⁵)
% of CD8
D
E
Day 7
Day 31
KLRG1+
CD43+ CD27+
CD43- CD27-
F
G
H
I
60%
35%
37%
53%
28%
17%
1.8%
47%
39%
12%
1.6%
50%
36%
12%
1.6%
KLRG1
CD127
CD27
CD43
% of GP33+
Cells per uL of Blood (10²)
Cells per spleen (10⁴)
Cells per spleen (10⁵)
miR-15/16fl/fl
Malat1Scr/Scr;miR-15/16Δ/Δ
miR-15/16Δ/Δ
